# Cross-linking/Mass Spectrometry: A Community-Wide, Comparative Study Towards Establishing Best Practice Guidelines

**DOI:** 10.1101/424697

**Authors:** Claudio Iacobucci, Christine Piotrowski, Ruedi Aebersold, Bruno C. Amaral, Philip Andrews, Christoph Borchers, Nicolas I. Brodie, James E. Bruce, Stéphane Chaignepain, Juan D. Chavez, Stéphane Claverol, Jürgen Cox, Gianluca Degliesposti, Meng-Qiu Dong, Nufar Edinger, Cecilia Emanuelsson, Marina Gay, Michael Götze, Fabio C. Gozzo, Craig Gutierrez, Caroline Haupt, Albert J. R. Heck, Franz Herzog, Lan Huang, Michael R. Hoopmann, Nir Kalisman, Oleg Klykov, Zdeněk Kukačka, Fan Liu, Michael J. MacCoss, Karl Mechtler, Ravit Mesika, Robert L. Moritz, Nagarjuna Nagaraj, Victor Nesati, Robert Ninnis, Petr Novák, Francis J O’Reilly, Matthias Pelzing, Evgeniy Petrotchenko, Lolita Piersimoni, Manolo Plasencia, Tara Pukala, Kasper D. Rand, Juri Rappsilber, Dana Reichmann, Caroline Sailer, Chris P. Sarnowski, Richard A. Scheltema, Carla Schmidt, David C. Schriemer, Yi Shi, J. Mark Skehel, Moriya Slavin, Frank Sobott, Victor Solis-Mezarino, Heike Stephanowitz, Florian Stengel, Christian E. Stieger, Michael Trnka, Marta Vilaseca, Rosa Viner, Yufei Xiang, Sule Yilmaz, Alex Zelter, Daniel Ziemianowicz, Alexander Leitner, Andrea Sinz

**Author notes:** Both authors contributed equally (Claudio Iacobucci, Christine Piotrowski).

## Abstract

The number of publications in the field of chemical cross-linking combined with mass spectrometry (XL-MS) to derive constraints for protein three-dimensional structure modeling and to probe protein-protein interactions has largely increased during the last years. As the technique is now becoming routine for *in vitro* and *in vivo* applications in proteomics and structural biology there is a pressing need to define protocols as well as data analysis and reporting formats that are generally accepted in the field and that have shown to lead to high-quality results. This first, community-based harmonization study on XL-MS is based on the results of 32 groups participating worldwide. The aim of this paper is to summarize the status quo of XL-MS and to compare and evaluate existing cross-linking strategies. From the results obtained, common protocols will be established. Our study serves as basis for establishing best practice guidelines in the field for conducting cross-linking experiments, performing data analysis, and reporting formats with the ultimate goal of assisting scientists to generate accurate and reproducible XL-MS results.

---

Mass spectrometry (MS) is becoming increasingly popular in the field of structural biology, with great implications for solving important biological questions. A central technique in structural MS is chemical cross-linking combined with MS (XL-MS). Since 2000, XL-MS and computational modeling has advanced from investigating three-dimensional structures of isolated proteins to deciphering protein interaction networks^1,2,3,4^. In the field of integrated structure analysis, XL-MS is often used in conjunction with cryo-electron microscopy. As the chemical XL-MS approach allows the capture of transient and weak interactions, it is now becoming a routine technique for unraveling protein interaction networks in their natural cellular environment^5^. The knowledge obtained will significantly advance our understanding of the structure of functional complexes, the topology of cellular networks and molecular details underlying human pathologies.

Briefly, the XL-MS approach relies on adding a chemical reagent to a protein solution connecting two functional groups of amino acid side chains. Cross-linker molecules consist of two reactive groups that are separated via a spacer of defined length and are often referred to as a “molecular rulers”. The cross-linked residues are usually identified after enzymatic digestion of the covalently connected protein(s) using LC/ESI-MS/MS (liquid chromatography electrospray ionization tandem mass spectrometry) and the resulting fragment ion spectra are computationally assigned to the cross-linked peptides. The distance constraints imposed by the chemical cross-linker on the protein’s tertiary structure serve as a basis for subsequent computational modeling studies to derive three-dimensional structural models. XL-MS can be applied to both proteins and protein complexes and in the case of protein assemblies, the distance constraints can be used to map the subunit topology. XL-MS is now increasingly being used for deriving protein-protein interaction maps, both *in vitro* and *in vivo*, where interacting proteins are covalently connected by the cross-linking reaction.^6,7,8,9,10^

The wide acceptance of XL-MS by the proteomics and structural biology communities reflects the increasing importance of cross-linking data for elucidating protein structures and protein-protein interactions. However, the growth of the user base brings about challenges of its own: Even a relatively superficial glance at the literature shows a huge diversity of cross-linkers, experimental workflows, and computational pipelines. Moreover, the information provided in scientific research articles that contain cross-linking data can range from being quite detailed to very brief.

The heterogeneity of cross-linking protocols has mainly emerged from the use of different cross-linking chemistries and different designs of the corresponding cross-linker (e.g., non-cleavable/cleavable, isotope-coded, or affinity-tagged reagents). This, in turn, necessitated individual software solutions specifically tailored to the analysis of data from the experimental workflow. The most common database search engines used in proteomics are not directly suitable for interpreting mass spectra from cross-linked peptides. Therefore, the majority of computational solutions have emerged from laboratories that pioneered the application of XL-MS and created tools specifically tailored for the analysis of cross-linked peptides. Together with a current lack of formal or even informal reporting standards, the present state of XL-MS may confuse researchers that are interested in adopting the technology. Currently, it is not clear, which strategies are most suitable in general or for a particular application, which makes it challenging to objectively compare results obtained by different groups.

Certainly, the challenges summarized above resemble those of other disciplines. In particular, scientists active in “conventional” proteomics research have tried to address the very same issues over the last decade. Inter-laboratory and software comparison studies have been performed for different experimental strategies, including data-dependent acquisition^11^, selected reaction monitoring^12, 13, 14, 15^, and most recently, data-independent acquisition^16, 17^. In addition, regular comparative studies have been organized by the Association of Biomolecular Resource Facilities (ABRF; https://abrf.org/research-group/proteomics-research-group-prg and https://abrf.org/research-group/proteomics-standards-research-group-sprg). Together, these studies revealed limitations in commonly used experimental and computational workflows, but on the other hand also provided evidence for the robustness of a particular technique when implemented in different laboratories according to standard operating procedures.

Standardized file formats and reporting guidelines for proteomics have been developed under the auspices of the Proteomics Standards Initiative (PSI) of the Human Proteome Organization (http://www.psidev.info)^18^. For example, as far back as 2007, the first recommendations for minimum reporting standards in proteomics (Minimum Information About a Proteomics Experiment, MIAPE) have been made^19^, which have been followed by detailed guidelines of several proteomics journals. PSI has also formalized open-file formats, such as the mzML format for raw MS data^20^ and the mzIdentML format for protein identifications^21^. Such guidelines and open data formats have also led to an increase in the deposition of proteomics data in open data repositories such as the PRoteomics IDEntifications (PRIDE) archive, hosted by the European Bioinformatics Institute (https://www.ebi.ac.uk/pride/archive/)^22^. via the ProteomeXchange initiative (https://www.proteomexchange.org)^23^.

Initiatives for structural MS applications, including XL-MS, ion mobility-MS, hydrogen/deuterium exchange, and native MS are only emerging. However, there is also a clear need for the objective assessment of the methods and reporting standards in these disciplines. For this purpose, several researchers active in the field of XL-MS decided to start a community-organized effort with the goal of providing a first overview of common procedures in XL-MS to generate the basis for best practices in the field.

In this first inter-laboratory effort 32 groups worldwide contributed, delivering a total of 58 cross-linking data sets. The data reflect the great diversity of experimental and computational strategies employed and to our knowledge, this is the first comprehensive study with the aim to harmonize the XL-MS field.

## Results

### Study Design

We opted for a simple study design to encourage participation from as many laboratories as possible, including those with currently only little experience in XL-MS. Invitations were sent out to research groups known to be active in the field from their published work and to attendants of the Symposium of Structural Proteomics (SSP, http://www.structuralproteomics.net/) meeting series. The guidelines were kept quite simple and each participant was provided with a template spreadsheet to document their method and report their results (Supporting Information). Bovine serum albumin (BSA), a protein with a molecular weight of ^~^66 kDa, was selected as study system. We requested that a certain product from a widely available supplier should be used, and it was specified to use a BSA concentration of 10 μM. Apart from these restrictions, we left the contributing labs full freedom to choose the experimental and computational strategies of their choice. This included, among other parameters, flexibility regarding the choice of cross-linking reagent and its concentration, buffer composition and pH, reaction time and temperature, postcross-linking sample processing (digestion protocol, optional fractionation and enrichment of cross-linked products), conditions for LC/MS analysis, and data analysis procedures (choice of software, search parameters, validation of the results). In short, we expected that participants would use the typical XL-MS workflows established in their labs. The protocols used by the individual participating labs were collected and analyzed in the Sinz lab and are summarized in the Supporting Information. For data analysis, we provided the amino acid sequence of mature BSA after cleavage of the signal peptide and propeptide sequences (residues 25-607 of the UniProt entry P02769; https://www.uniprot.org/uniprot/P02769) to ensure a uniform numbering scheme. Finally, we encouraged participants to perform at least three replicates. As mentioned above, we provided a template spreadsheet (Supporting Information) that needed to be completed by the participants before a data set would be considered for inclusion in the detailed assessment of the results. An overview of the data sets provided by different labs is presented in Figure 1.

**Figure 1:**
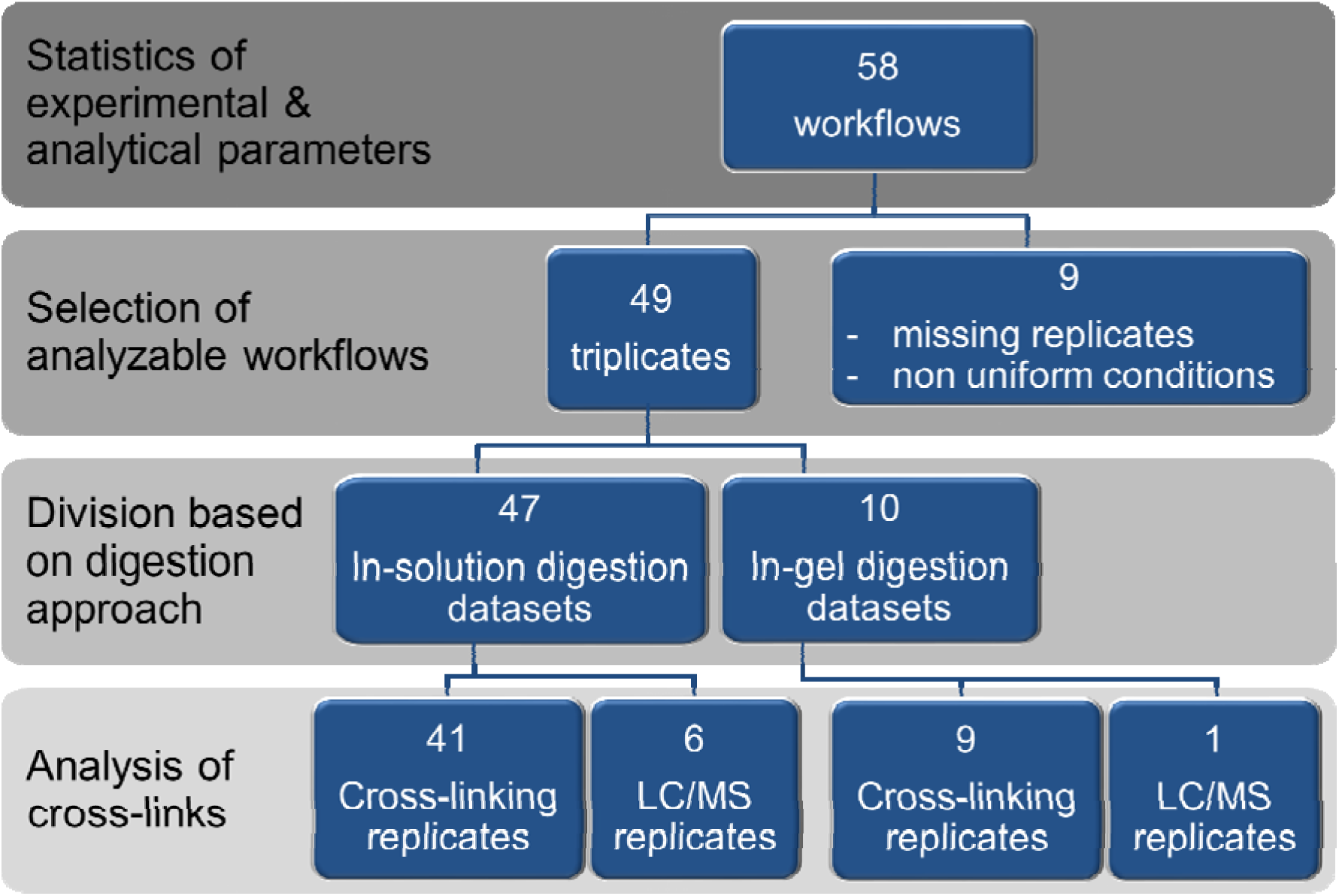
Overview of data sets provided by the participants of this study. 32 groups participated in this study, yielding 58 separate cross-linking workflows. Nine datasets had to be excluded due to missing replicates and non-uniform conditions, resulting in a total of 49 datasets that were further considered. Several workflows contain both in-solution (47 samples) as well as in-gel digestion (10 samples) as processing methods. The samples were considered only once during a workflow analysis.

### Protein System

BSA was selected as model protein for this study as it is a globular and stable protein that is readily available at low cost. Moreover, the three-dimensional structure of BSA is well-known and we selected the Protein Data Bank entry 4F5S (https://www.rcsb.org/structure/4F5S) for further interpretation of the results. As BSA possesses a tendency towards forming dimers, this has to be considered when interpreting the results (see also below).

### Cross-linking Reagents

As outlined above, the participants of this study were free to choose the cross-linking principle(s) on their own. The majority of groups decided to use non-cleavable, homobifunctional, amine-reactive *N*-hydroxysuccinimide (NHS) cross-linkers, *i.e.*, bis(sulfosuccinimidyl)suberate (BS^3^) or disuccinimidylsuberate (DSS) (Figure 2A). Both cross-linkers only differ by a sulfonic acid group that is incorporated for increased water solubility and bridge a distance of 11.4 Å, resulting in C_α_-C_α_ distances of ^~^27 Å to be cross-linked^24^. MS-cleavable cross-linkers, such as disuccinimidylsulfoxide (DSSO) and disuccinimidyldibutyric urea (DSBU), are increasingly being used as they allow a targeted identification of cross-linked product based on characteristic reporter ions generated during MS/MS experiments. MS-cleavability as a cross-linker feature is essential to reduce the search space in conducting proteome-wide cross-linking studies. The vast majority of cross-linkers used herein target amine groups in proteins, i.e., lysine side chains, while carboxylic acid groups, such as aspartic and glutamic acid residues, are less frequently targeted (Figure 2B). The main spacer lengths of the cross-linkers are determined by the three most abundant cross-linkers used in this study are: BS^3^ and DSS (both 11.4 Å), DSBU (12.5 Å), and DSSO (10.1 Å) (Figure 2C).

**Figure 2:**
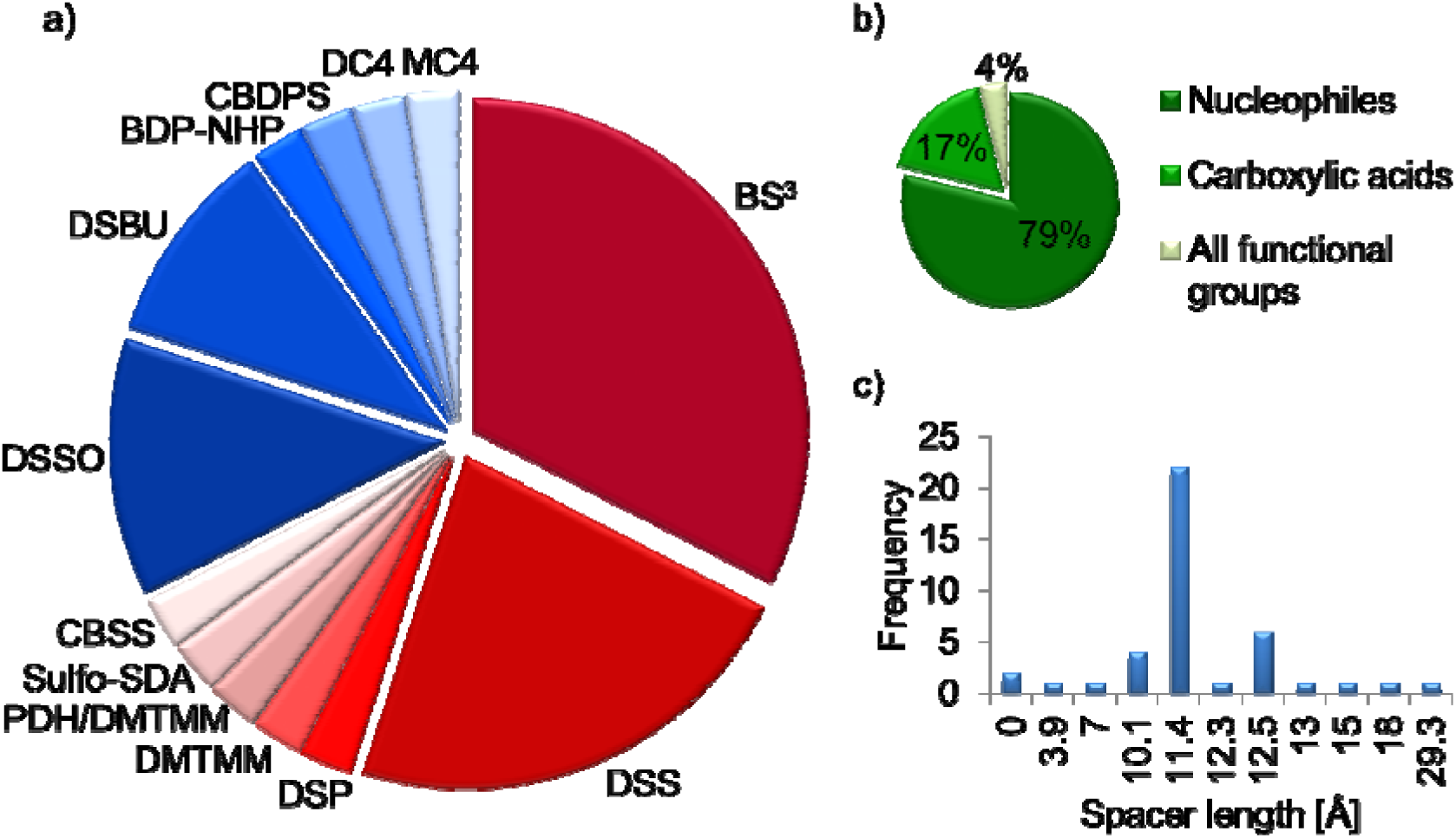
(A) Cross-linking reagents used in this study, (B) reactivity, (C) spacer length. The cross-linkers used in this study are: BS^3^ (bis(sulfosuccinimidyl)suberate, DSS (disuccinimidylsuberate), DSP (dithiobis(succinimidylpropionate)), DMTMIVI (4-(4,6-dimethoxy-1,3,5-triazin-2-yl)-4-methyl-morpholinium chloride) with and without PDH (pimelic acid dihydrazide), sulfo-SDA (sulfosuccinimidyl 4,4’-azipentanoate), CBSS (carboxybenzophenone sulfosuccinimide), DSSO (disuccinimidylsulfoxide), DSBU (disuccinimidyldibutyric urea), BDP-NHP (*N*-hydroxyphthalamide ester of biotin aspartate proline), CBDPS (cyanurbiotindimercaptopropionyl succinimide), DC4 (1,4-bis(4-((2,5-dioxopyrrolidin-1-yl)oxy)-4-oxobutyl)-1,4-diazabicyclo[2.2.2]octane-1,4-diium), and MC4 (N,N’-bis(4-((2,5-dioxopyrrolidin-1-yl)oxy)-4-oxobutyl (-morpholine).

### Reaction Conditions

The reaction conditions were also kept completely open to the participants, including cross-linking reaction time, temperature, cross-linker excess, and pH value of the cross-linking solution (Figure 3). Not surprisingly, the pH value of the cross-linking reaction mixture was kept around pH 7.4 to 7.5 in the majority of experiments in order to resemble the physiological pH situation. A pH value of 8.0 that was also used in some experiments has the advantage of enhancing the reactivity of NHS esters with nucleophiles. The temperature was kept to 20, 25 or 37 °C in the majority of experiments, with lower temperature being applied only by a few groups. For BSA, a temperature of 37 °C certainly does not present a problem as it is a stable, globular protein, but for delicate and unstable proteins one should take care to conduct the cross-linking reaction at lower temperatures.

**Figure 3:**
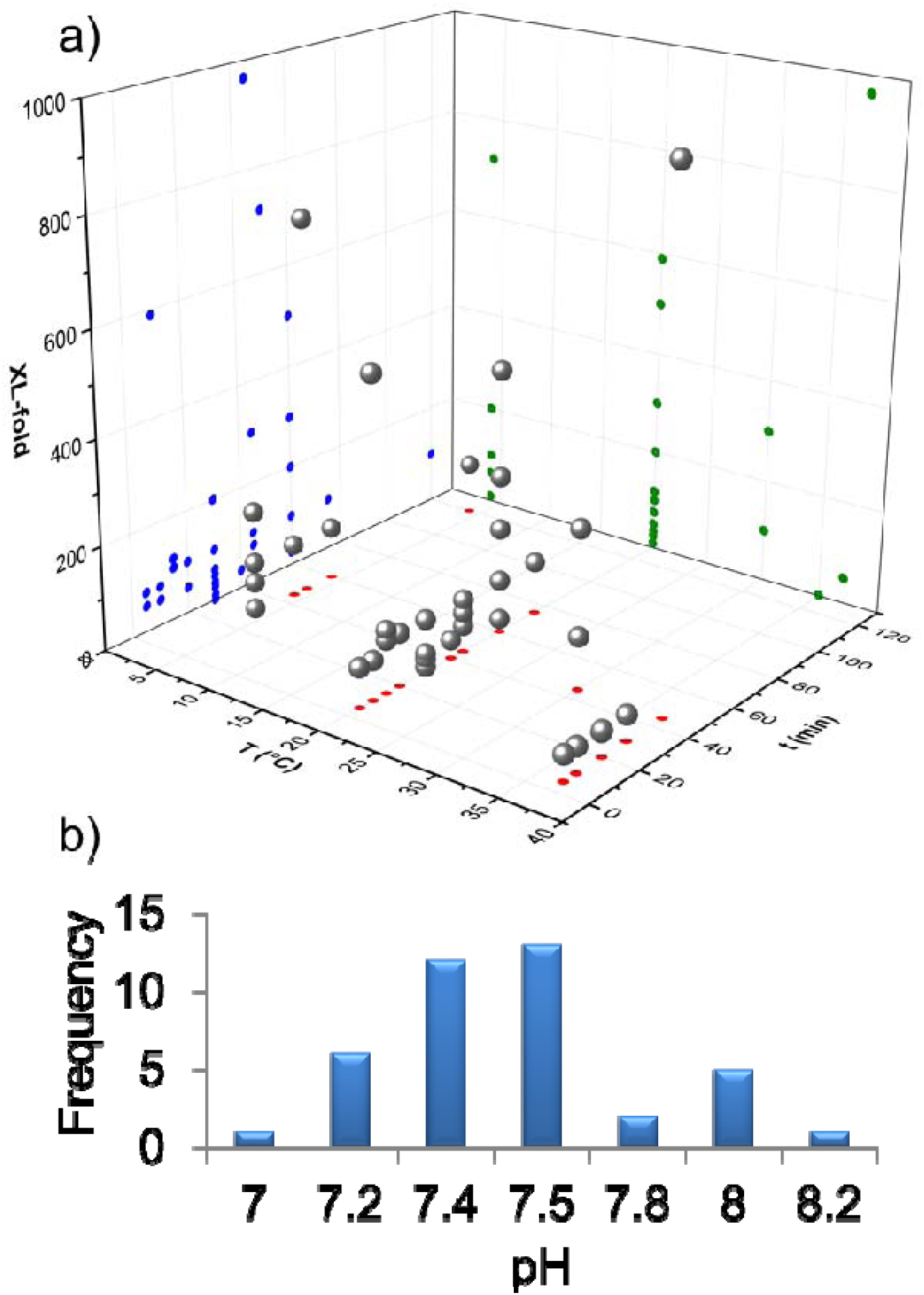
Cross-linking reaction conditions. (A) Time, temperature, and cross-linker excess (XL-fold) as well as (B) pH values of the cross-linking reactions were variable parameters.

### Instrument Platforms and Settings Used to Generate XL-MS Data

The overwhelming majority of cross-linking data were generated on Orbitrap mass spectrometers (Figure 4). Only two FTICR (SolariX and Velos FTICR) mass spectrometers and one Q-TOF (Synapt G2 SI) instrument were employed (Figure 4A). All groups used LC/ESI-MS/MS analysis, applying for most experiments a resolving power of 60,000 or 120,000 (at *m/z* 200 or 400, as specified by the manufacturer Thermo Fisher Scientific for orbitrap instruments) (Figure 4B). For MS/MS experiments, a resolving power of 15,000 or 30,000 was employed in most cases (Figure 4C). Details on enrichment of cross-linked species, considered charge states, fragmentation methods, and MS^3^ resolution are presented in the Supporting Information (Figure S1).

**Figure 4:**
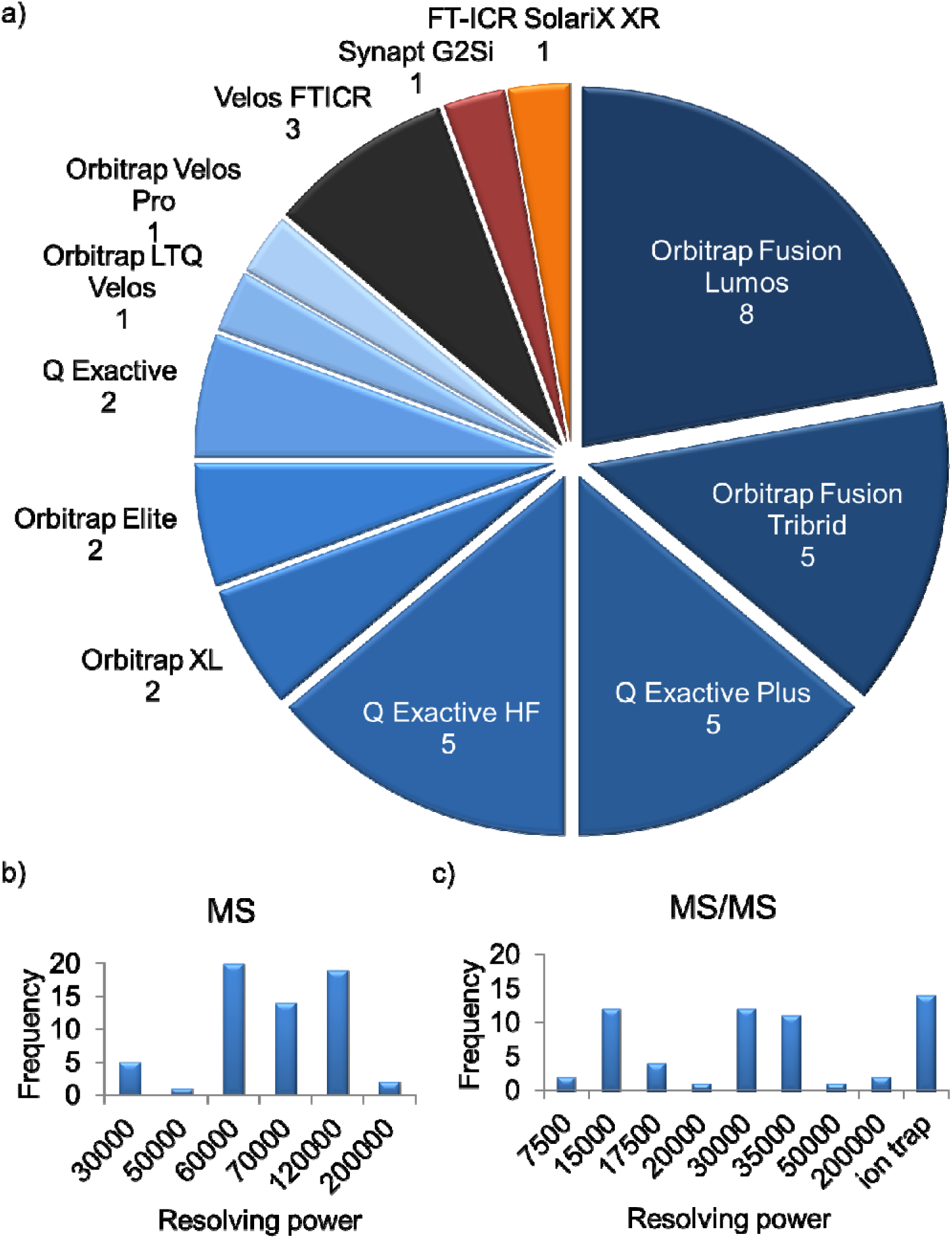
LC/MS/MS conditions applied. (A) MS instrumentation, (B) MS resolving power, (C) MS/MS resolving power. Resolving power is defined at *m/z* 200 for orbitrap instruments, while for ICR instruments it is defined at *m/z* 400. Please note that several research groups generated datasets with different instruments and settings.

### Data Analysis and Validation Strategies

Strategies for data analysis were highly diverse (Figure 5), reflecting the variety in the XL-MS field where nearly every group possesses their own software tools tailored to fit their specific needs. This enormous variety is currently one of the most critical issues in XL-MS and we consider it as important contribution of this study to reflect this diversity. The false discovery rate (FDR) plays an important role in this context and from this study it arose that most of the groups apply an FDR of 5% (Figure 5B). Manual validation of the cross-links was performed for 66% of the experiments, while in 34%, the data sets were not manually checked. It is important to note that a mechanism to control the FDR should exist in the software, although proper FDR control is not trivial for small search spaces, manual validation strategies might be especially beneficial in such cases. Some strategies provide additional layers of evidence that can be used to better control the error rate. For example, isotope-coded, non-cleavable linkers provide two independent measures of precursor and fragment masses, and charge state information for fragments independent of MS resolution; MS-cleavable linkers provide three layers of information - intact precursors, released fragments corresponding to intact peptide chains, and fragments thereof. In absence of such strategies, we recommend that preferentially both, MS and MS/MS data, should be recorded with high mass accuracy to rule out a false assignment of cross-linked products. Clearly, some of these effects will only become apparent for samples of higher complexity.

**Figure 5:**
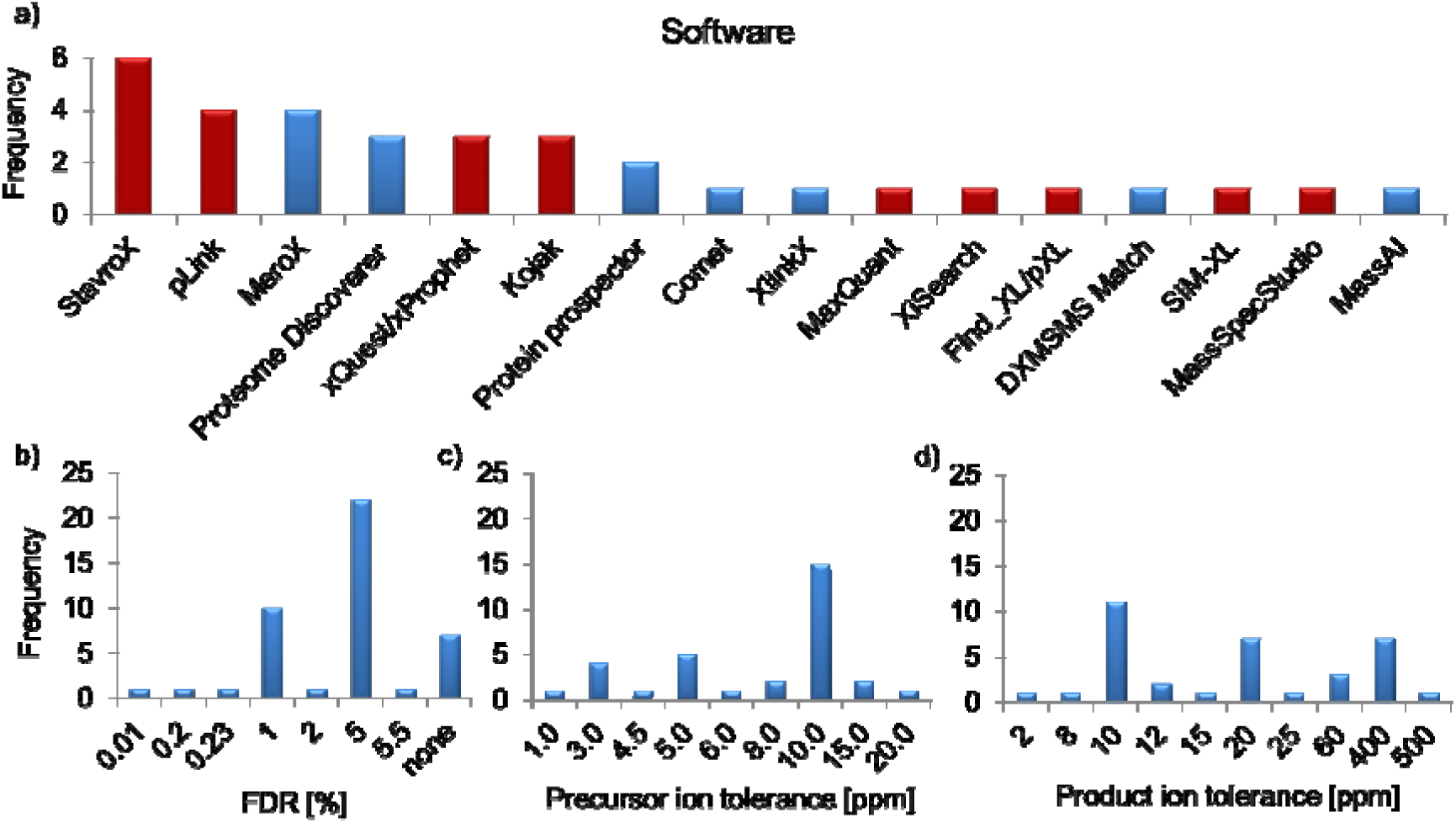
(A) Software tools used in this study. Red bars indicate that the software is applicable only for non-cleavable cross-linkers; blue bars indicate that the software can be used for MS-cleavable cross-linkers. (B) False discovery rates. (C) Mass tolerances MS. (D) Mass tolerances MS/MS. For the Proteome Discoverer, data analysis was performed using the XlinkX software node.

### Identified Cross-links

As we left it to the individual participants whether to use in-solution or in-gel digestion as work-up method before LC/MS/MS analysis, 47 data sets were generated by in-solution digestion, while 10 samples originated from in-gel digestion (Figure 1). As already mentioned, BSA has a tendency to form dimers, which somewhat complicates data analysis. In case only the BSA monomer band is used for in-gel digestion and subsequent generation of the cross-linking data set, one can definitely rule out that cross-links are in fact representing intermolecular interactions between two BSA molecules. On the other hand, during the in-gel digestion procedure cross-links might get lost, resulting in an overall lower number of cross-linked products.

Another aspect regards the reaction sites that were considered during data analysis. Usually, NHS esters, such as the mainly used cross-linkers BS^3^, DSS, DSBU, and DSSO will react with lysine, but they also exhibit a significant reactivity towards serine, threonine and tyrosine. The pH used for conducting the cross-linking reaction plays a significant role as amine reactivity is increased at higher pH values. Some participants considered only Lys-Lys cross-links and neglected the side-reactivity of NHS esters with hydroxy group-containing amino acids. In this study, it became apparent that Ser, Thr, and Tyr account for ^~^30% of cross-linking sites (Supporting Information, Figure S2). The reactivity of Ser, Thr, and Tyr residues obviously depends on the reaction conditions (cross-linker, pH value of the solution) as well as local pK_a_ value. It is not practicable to consider Lys, Ser, Thr, and Tyr when analyzing very complex systems, such as complete proteomes. Therefore we suggest as a compromise to consider for whole proteome samples only lysine as reactive sites of NHS ester cross-linkers, while for single proteins or proteins assemblies, Lys, Ser, Thr, and Tyr might be taken into account.

Figure 6 provides an overview about the reproducibility of results obtained with the individual workflows of the participants. For in-solution digestion workflows, the average number of unique cross-links in BSA is 85.7, while for in-gel digestion workflows using only the monomeric BSA band, the average number is 44. The term “cross-link” refers to the specific amino acid residues that are connected, irrespective of different peptide sequences due to missed cleavage sites or modifications. The majority of participating labs came up with similar numbers of unique cross-links, independently of the cross-linking conditions used (Figure 6A). Three cross-linking workflows however recorded a significantly higher number of cross-links (between 260 and 350). The reason could be a false consideration of cross-links from BSA dimers that in some preparations might have been a dominating species due to inappropriate sample treatment. For in-gel digestion workflows, up to 19 overlength cross-links were reported in one dataset, which could represent false-positives due to partial unfolding as only the monomeric form of BSA was considered in these samples (Figure 6B).

**Figure 6:**
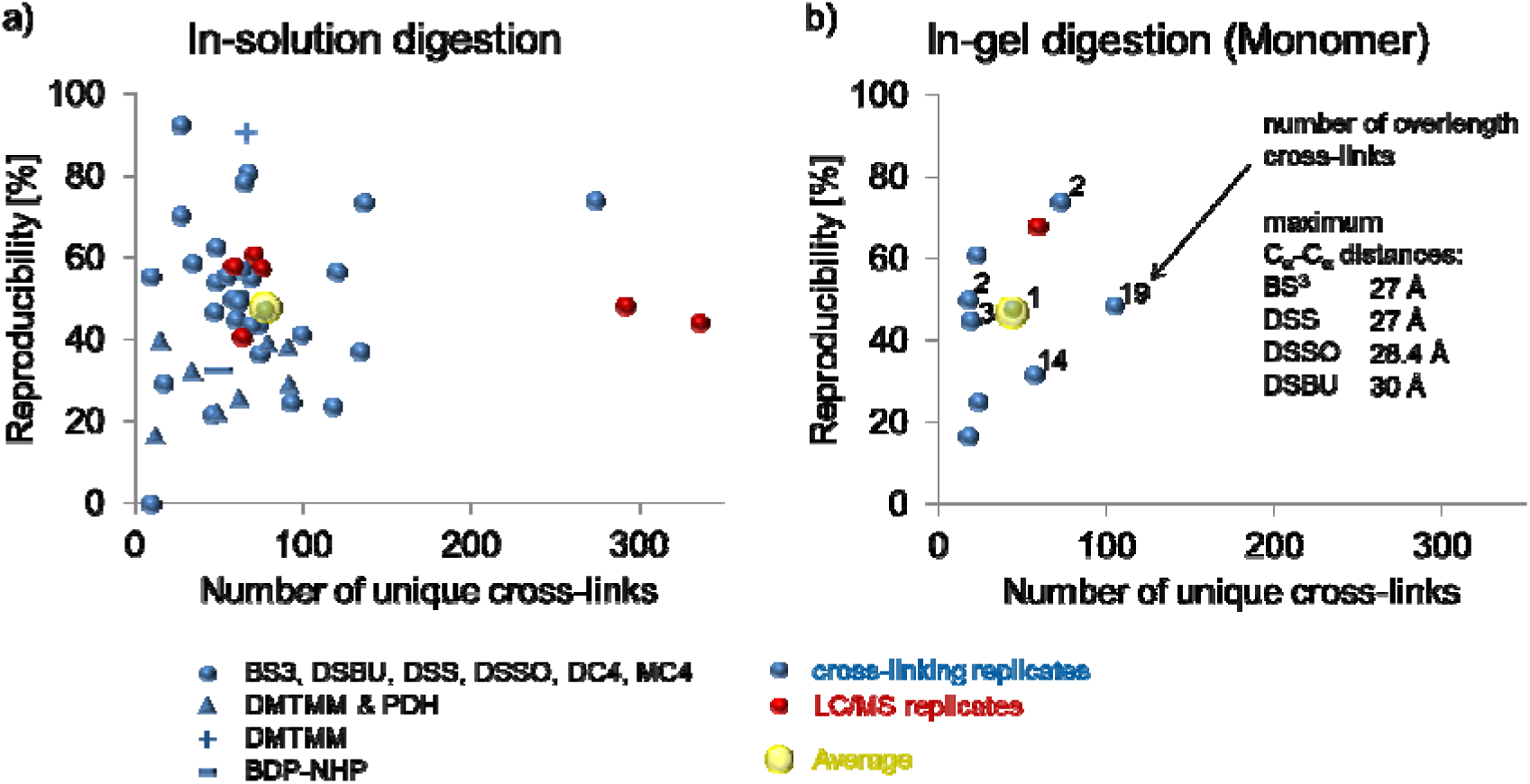
Number of BSA cross-links identified. (A) in-solution, (B) in-gel digestion workflows.

A more detailed inspection of the unique cross-links revealed highly interesting insights: Datasets created from amine-reactive cross-linkers (BS^3^, DSBU, DSS, DSSO, DC4, MC4, CBDPS) using an in-solution digestion workflow yielded a total of 1066 unique cross-links. A complete list of unique cross-links, identified with cross-linkers reacting with nucleophiles (amine and hydrox groups) and sorted by their reproducibility, is provided as separate file in the Supporting Information. 601 of 1066 unique cross links (56%) were however identified in only one single dataset (Figure 7). This indicates an overall low reproducibility of cross-linking results. The curve in Figure 7A shows that the number of unique cross-links identified is inversely proportional to the reproducibility of cross-links in the data sets (coefficient of proportionality ≃ −1). If the reproducibility across the data sets is higher than 20%, the effect of including more datasets, different reaction conditions, and analytical parameters determines a linear increment of the number of cross-link identifications. The intercept with the y-axis of the resulting interpolated linear curves indicates the putative ideal number of cross-links in BSA to be between 73 and 88 (Figures 7B and 7C). This value is very close to the average number of cross-links found (85.7 cross-links per dataset for in-solution digestion workflows, Figure 6A).

**Figure 7:**
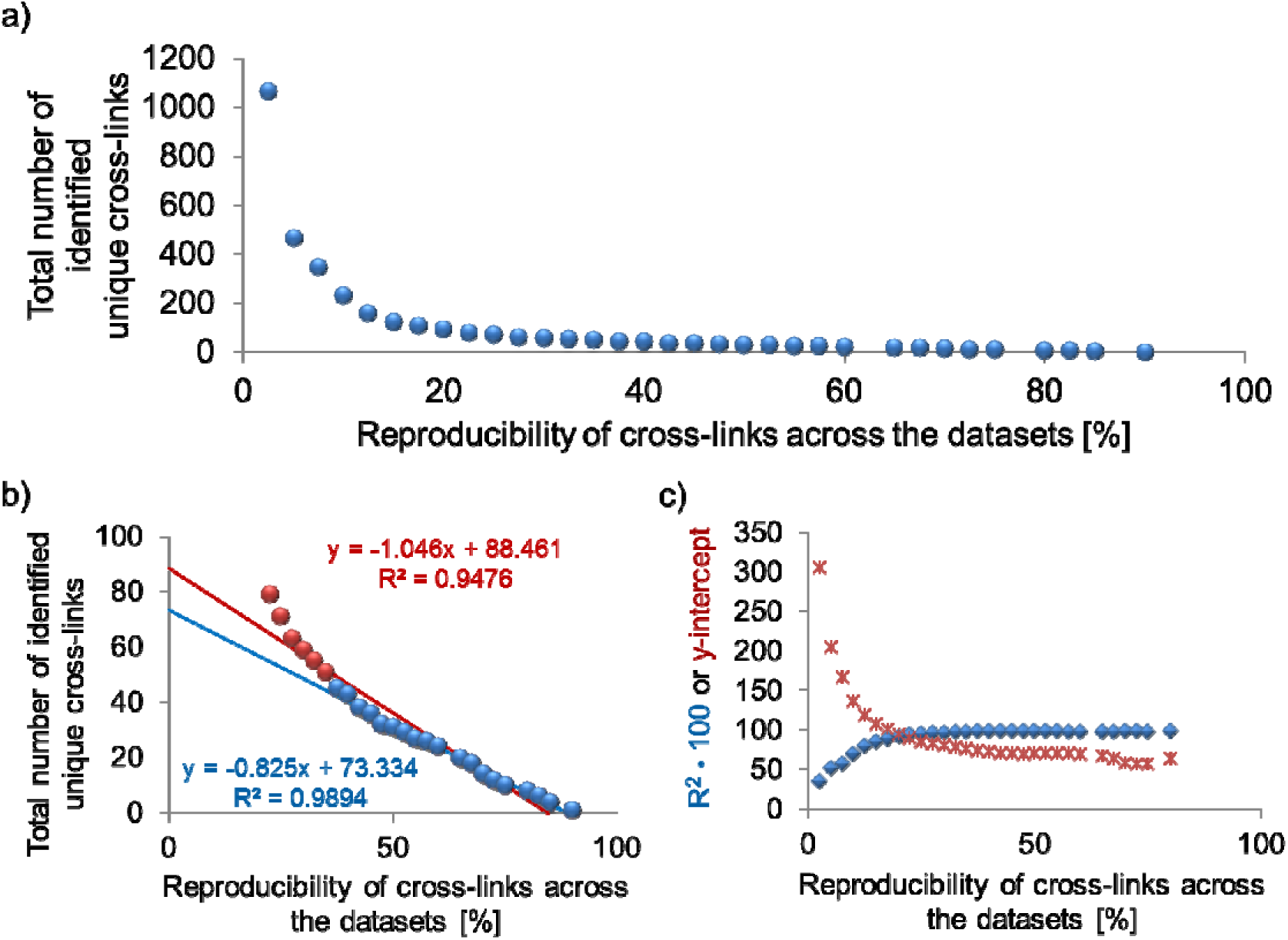
Comparison of unique cross-links (A, B, C). “Cross-link” defines the amino acid residues that are connected.

### Comparison of data acquisition and analysis strategies from one participating lab

Because most of the data in this study have been generated in different laboratories, differences in instrumentation and in the software used for data analysis make a direct comparison of selected results difficult. However, we used a subset of the data generated in a single laboratory to study the effect of the type of mass spectrometer and of different search settings on the outcome for a relatively simple model system, such as BSA (see Supporting Information).

## Discussion

This first community-based cross-linking study reflects the high diversity of XL-MS workflows that are currently employed in different labs worldwide. However, it also became apparent that independently of the workflow used the results obtained are to some degree comparable. For beginners in the field we suggest to use BSA as an initial study system and compare the outcome to the results obtained herein. As a guideline, the number of cross-links expected for BSA should be ca. 88 for an in-solution workflow, considering cross-links of the monomer and the dimer. Not unexpectedly, our study did not reveal *the* optimum experimental protocol or software to be used in any and all projects. The applications of XL-MS are just too diverse so that no single cross-linker, instrument or software tool is expected to be preferable for all scenarios, ranging from single protein (as used in this work) to whole-cell cross-linking. There are also clear interdependencies between the type of cross-linker (cleavable, non-cleavable) and the software that can be applied to process such data, as well as between instrument type and software as not all fragmentation methods or other MS platform-dependent features may be supported.

As discussed above, XL-MS has become an essential part of many structural proteomics studies, but is also a key element in integrative structural biology projects. In such interdisciplinary work, XL data may only be a small “puzzle piece” that is combined with other experimental data provided by methods such as electron microscopy, X-ray crystallography, NMR spectroscopy, small-angle X-ray scattering, together with computational modeling. Details about how experiments were carried out, how the data were processed, and how error rates were assessed are often missing from the publication, making it difficult for reviewers and readers to assess the reliability and credibility of the results. We therefore recommend that appropriate consideration should be given to the method section of all XL-MS publications by providing all necessary experimental and computational details. Our reporting template could serve as a starting point for the “minimum information about a cross-linking experiment” that should be included in research articles containing XL-MS data. Sufficient information needs to be provided, irrespective of the relative contribution of the cross-linking experiments to a specific project. This will also facilitate the cross-referencing of XL-MS data in integrative structural biology projects, for example in the dedicated PDB prototype archive, PDB-Dev^25^.

Data deposition to a proteomics repository, such as PRIDE, is encouraged, as the paucity of available data sets do not assist the field in validation, methods evaluation and workflow quality. It should be noted that not all data sets assigned to the cross-linking category in PRIDE originate from genuine XL-MS experiments (in the sense that cross-linking sites were identified), but also contain data from experiments that used cross-linking for the stabilization of complexes. The low uptake of data deposition may in part be due to the specific nature of XL-MS data. For a “complete” submission to ProteomeXchange, allowing a complete integration of search results and assignment of a Digital Object Identifier, the reported results need to be compliant with a PSI format, such as mzIdentML. Although the most recent version of mzIdentML (version 1.2) includes support for some XL-MS strategies, such a proteomics-centered format cannot easily consider all possible workflows, and few dedicated cross-linking search engines offer mzIdentML-compliant export at this point. Nevertheless, even a “partial” submission will make the raw MS data and results available in a user-specified format for download and re-use by interested researchers.

Additional studies that cover a wider range of sample types, such as large multi-protein assemblies or even whole proteomes, will be required to obtain a better understanding of the benefits and drawbacks of different experimental workflows. However, we believe that this first community-based study serves as the starting point for further initiatives in this direction, and encourages the adoption of consistent reporting and data sharing guidelines in XL-MS. We would like to invite interested parties to participate in the discussion to expand the community.

## Methods

All workflows employed by the participating groups are presented in a spreadsheet to document their methods and to report their results (Supporting Information).

## Acknowledgements

This study was conducted within the EU COST Action BM1403.

## Author contributions

CI, CP, AL, AS designed the study and wrote the manuscript, all authors provided data.

The authors do not have a conflict of interest.

## Additional information

Content of Supporting Information:

**Figure S1:** Details on enrichment of cross-linked species, considered charge states, fragmentation methods, and MS^3^ resolution.

**Figure S2:** Influence of cross-linking sites considered in data analysis.

**Figure S3:** Comparison of non-redundant BSA cross-links identified on three different orbitrap MS platforms.

**Figure S4:** Comparison of different data analysis strategies for DSS data sets acquired on Orbitrap Elite or Orbitrap Fusion Lumos instruments.

List of unique cross-links (pdf file).

Data reporting template (Excel file).

Data provided by all participants of this study (zip file).

